# The respiratory syncytial virus surge in Austria, 2022, was caused by lineages present before the COVID-19 pandemic

**DOI:** 10.1101/2023.01.26.525650

**Authors:** Monika Redlberger-Fritz, David N. Springer, Stephan W. Aberle, Jeremy V. Camp, Judith H. Aberle

## Abstract

In 2022, Austria experienced a severe respiratory syncytial virus (RSV) epidemic with an earlier-than-usual start and increased numbers of paediatric patients in emergency departments. Nationwide multiyear genomic surveillance revealed that the surge was driven by RSV-B, however genotypes consisted of multiple lineages that were circulating prior to the pandemic.

---

In 2022, a severe RSV epidemic occurred in Austria and in other countries [1,2] with an unprecedented burden of paediatric patients in emergency departments [3]. The surge began in the autumn, occurring earlier than in previous seasons, with the exception of the first RSV season following the emergence of COVID-19, which began in the summer 2021. The implementation of COVID-19 non-pharmaceutical interventions (NPI) in many countries, including Austria, resulted in a strong reduction of RSV activity during winter 2020/21. Recent studies showed that this prolonged lack of virus exposure resulted in a decline of RSV-specific immunity [4,5], which may explain the earlier-than-usual start in 2021, but may not fully explain the drastic increase in cases in 2022. An alternative or potentially additional hypothesis to explain the surge could be the emergence of novel genotypes with increased transmission or pathogenicity [6]. Therefore, we were interested in analysing the molecular epidemiology of the recent epidemics to better understand the phylodynamics of RSV. Here, we present data from RSV surveillance from 2012-2022, including an analysis of genomes from before and during the COVID-19 pandemic.

## Respiratory syncytial virus surveillance and genotyping in Austria

We tested 26,293 nasopharyngeal samples from the nationwide respiratory surveillance network that were collected year-round from ambulatory and hospitalized patients (median age 29yr, range 0-102yr) at 248 locations between calendar weeks 40/2012 and 47/2022. The 2021 and 2022 RSV epidemics started in calendar weeks 35 and 45, much earlier than observed in the pre-pandemic years 2012 to 2019, which followed a typical seasonal pattern with onset in early winter, between weeks 48 and 2 (**Figure 1a**). In comparison to 2019 and 2021 epidemics, which were dominated by the RSV-A subtype, the 2022 surge in Austria was driven by RSV subtype B strains (**Figure 1a**).

**Figure 1.**
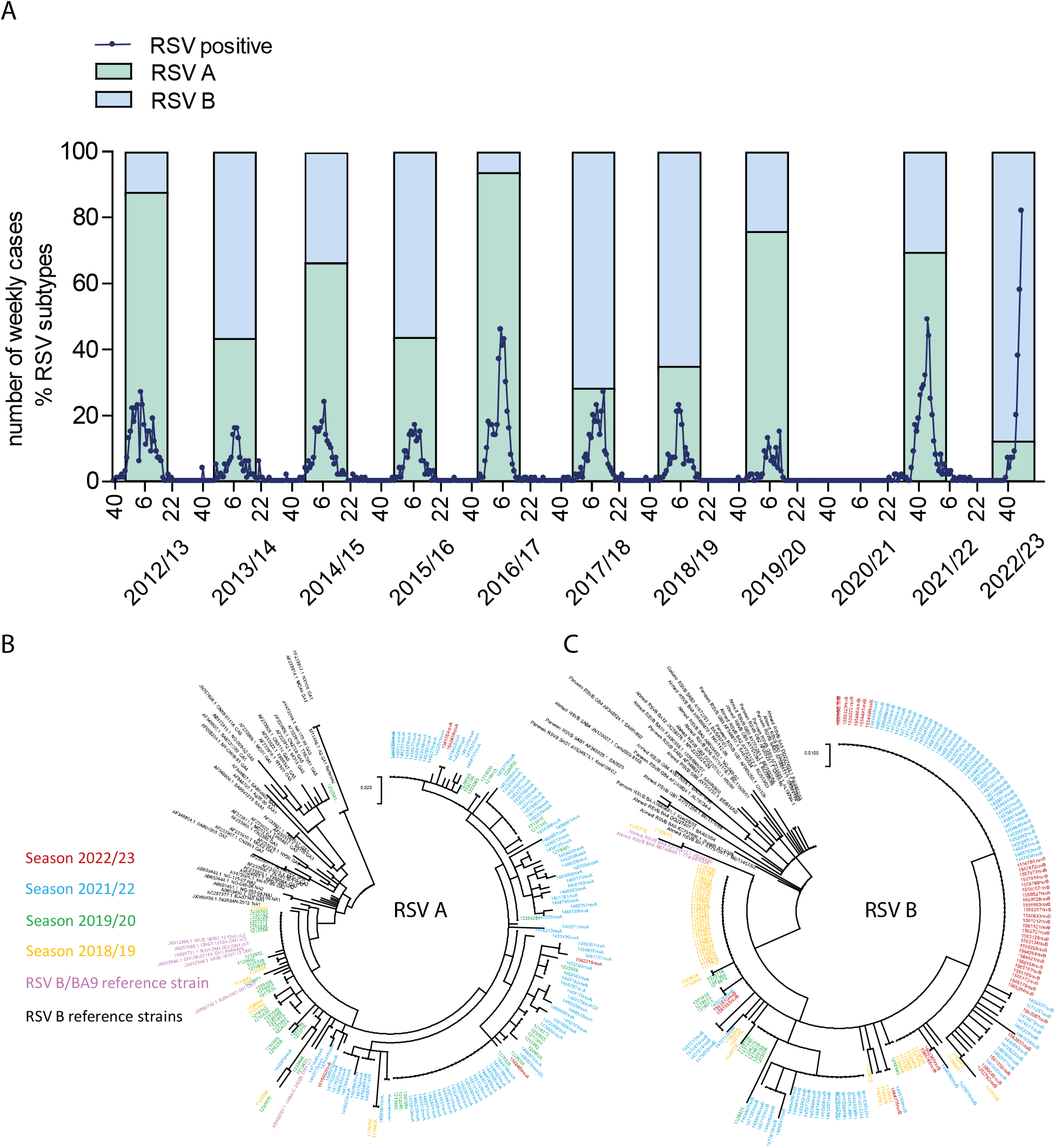
Epidemiological and phylogenetic analysis of respiratory syncytial virus strains from 2012 to 2022 in Austria. (A) Weekly Respiratory syncytial virus (RSV) surveillance during 2012 to 2022 by the nationwide respiratory surveillance network in Austria, as indicated by the number of RT-PCR positive samples for RSV (blue line). Stacked bars represent percentages of RSV groups A (green) and B (blue). (B) Phylogenetic analysis of RSV B genomes (n=187) and (C) RSV A genomes (n=186) using the maximum likelihood method based on the Tamura-Nei model of a partial sequence (236 nt) from the G protein. Colours indicate the seasons in which sequences were detected: 2022 in red, 2021 in blue, 2019/20 in green, 2018/19 in yellow, RSV B/BA9 and A/ON1 reference strains in pink in panels (B) and (C), respectively, and historical reference strains in black.

## Respiratory syncytial virus whole genome sequencing and phylogenetic analysis

We performed Sanger sequencing of the RSV surface glycoprotein G second hypervariable region (236 nt) to evaluate changes in the RSV groups and genotypes circulating in 2021 and 2022, and the two seasons preceding the COVID-19 pandemic. Phylogenetic analyses, comparing these sequences to publicly available reference sequences (**Supplemental**), demonstrated that all 187 RSV-B sequences obtained from 2018 to 2022 were RSV genotype GB5.0.5a (BA9) (**Figure 1b**). Similarly, RSV-A genotype GA2.3.5 (ON1), identified in all 186 RSV-A sequences in 2021 and 2022, was detected in pre-pandemic years (**Figure 1c**).

Whole genome sequencing of eight RSV-A samples from this cohort (including three from 2022) and 13 RSV-B samples (seven from 2022) confirmed that the genotypes circulating in the current surge were present prior to and during the COVID-19 pandemic (**Figure S1; Supplemental**). Analysis of the genetic distances (p-distance) and phylogenetic distances (patristic distance) indicated that the intragenotypic diversity did not exceed the threshold for characterizing a new lineage (RSV-B p-distance = 0.007 [S.E. 0.0003], patristic distance = 0.008 [SE 0.0032]; RSV-A p-distance = 0.032 [SE 0.0003], patristic distance 0.014 [SE 0.00006]) [7,8]. However, samples from each subtype appeared to form distinct (sub)lineages with strong bootstrap support and shared mutations (**Figures S2 and S3**). For RSV-B, four samples from Austria and elsewhere in 2022 shared mutations in the G attachment glycoprotein (S100G/D, P216S, P223L, Y267H), the fusion glycoprotein (R42K, S190N, S389P), the M2 protein (D38N), and the L protein (K570R, V1479A, R1759K, V1965I). For RSV-A, at least three sublineages were present, one with G: L142S and L: N143D; one with G: A57V, G: K209R, G: L286P, G: T295I, and F: A103T; and one with L: L835M. A mutation at site G: T296A was shared by two of these sublineages. All sequences were submitted to GISAID and GenBank sequence databases (**Supplemental**).

## Discussion

Since 2020, the COVID-19 pandemic and implementation of NPIs have had broad effects on the spread of RSV in the community. Through the nationwide respiratory surveillance network, we have monitored changes in the circulation patterns of RSV year-round in Austria.

In the pre-pandemic years 2012 to 2019, RSV epidemics followed a typical seasonal pattern, with onset in early winter. With the implementation of NPIs early in the COVID-19 pandemic, there was a significant disruption of the typical circulation pattern with an almost complete absence of RSV cases during the 2020/21 season. The changes in community circulation have led to a decline of population-level immunity [4,5], which may explain in part the rise in RSV cases observed in 2021 and in 2022. Other factors, such as specific changes in the circulating genotypes and/or the emergence of a novel genotype, may be contributing to the increased spread [9].

One change observed in Austria is a shift in the RSV subtype predominance, with RSV-A genotype GA2.3.5 predominating in 2021, and RSV-B genotype GB5.0.5a predominating the surge in 2022. Phylogenetic comparisons of the complete RSV-A and RSV-B genomes indicate that the genotypes driving the 2022 epidemic in Austria had been circulating in pre-pandemic years. There was a notable expansion of an RSV-B sublineage with distinct differences compared to strains detected in the pre-pandemic seasons. While this sublineage appears to be the only RSV-B lineage circulating in the US, where the late 2022 season has been predominated by RSV-A, we detected additional RSV-B sequences more similar to pre-pandemic sublineages as well [6]. RSV-A genotype GA2.3.5 was notably more diverse than RSV-B, and several well-supported sublineages were represented in 2022 as well as in the four prior seasons. Together, the findings indicate that the RSV-A and -B lineages persisted throughout the pandemic and expanded in association with the surge of cases in 2021 and 2022. It remains to be determined whether the shared derived mutations in the identified sublineages, particularly in the newly emerged RSV-B sublineage, had a part in driving the unexpected surge in 2022.

The findings demonstrate the need for ongoing surveillance and monitoring of RSV genotypes, as recommended by the European Centre for Disease Prevention and Control [10]. Molecular epidemiology will be particularly helpful in understanding how RSV activity will change over the next years, in terms of seasonality, subtype circulation and lineage distribution, in the wake of the COVID-19 pandemic.

## Acknowledgments

The authors wish to thank Claudia Seeber for technical support.

## Declaration of interest

We declare no competing interests.

## Notes

### Competing Interest Statement

The authors have declared no competing interest.

