## Supplemental for "The respiratory syncytial virus surge in Austria, 2022, was caused by lineages present before the COVID-19 pandemic"

### Supplementary 1

##### Whole genome sequencing methods

Whole genome sequencing was performed with a tiled amplicon approach (Wang et al. 2022), using two primer pools specific to each subtype, RSV-A and RSV-B. Amplicons were pooled and a multiplexed library was prepared for sequencing using the Nextera XT kit (Illumina). Sequencing was performed on a MiSeq with V2 chemistry using 150 bp paired end reads. The reads were quality trimmed using fastp (v 0.23.2) and contigs were assembled using SPAdes (v 3.15.4). Reads were mapped to the contigs using bwa-mem2 (v 2.2.1), and consensus sequences were called using ivar (v 1.3.1). Sequences were submitted to both GISAID and GenBank databases.

##### Whole and partial genome phylogenetic analysis methods

Sequences were aligned to reference sequences obtained from GenBank public database using clustal omega (v 2.1). Phylogenetic analysis of whole genome sequences was performed with phyml (v 3.3.20211231), using the GTR+G substitution model, and inferring support over 500 bootstraps. To incorporate even more contemporary sequences where whole genomes were not available, phylogenetic analysis of full G protein open reading frame was performed in MEGA 7 with the GTR+G substitution model over 500 bootstraps. A full list of reference sequence accession numbers are provided below. Patristic distances were calculated using the “ape” package (v 5.6-2) in R (v 4.0.3), and within-genotype p-distances were calculated in MEGA 7 using bootstrap technique to calculate standard error. Phylogenies were drawn and annotated in ggtree (v 3.4.4; Yu, et al. 2017) with associated packages in R.

*List of accession numbers for reference sequences for RSV-A*

AY911262, JF920053, JF920062, JQ901452, JQ901455, JX015480, JX015482, JX015486, JX015491, JX015497, JX069798, JX069801, JX069802, KC731483, KF530260, KF826826, KF826827, KF826832, KF826838, KF826847, KF826850, KF826854, KF826855, KF973319, KF973333, KJ627256, KJ627284, KJ627294, KJ627305, KJ627320, KJ627336, KJ627337, KJ627338, KJ627349, KJ627370, KJ627647, KJ627655, KJ627656, KJ627717, KJ627719, KJ627733, KJ641590, KJ643463, KJ643464, KJ643589, KJ672455, KJ672467, KJ672470, KJ672482, KJ723465, KJ723474, KJ723483, KJ723486, KJ723492, KJ939951, KM360090, KP119748, KP258699, KP258700, KP258701, KP258709, KP258715, KP258723, KP258726, KP258743, KP317953, KP663728, KT285064, KU316092, KU316098, KU316104, KU316118, KU316133, KU316137, KU316149, KU316161, KU316165, KU316170, KU950506, KU950531, KU950540, KU950550, KU950556, KU950560, KU950573, KU950596, KU950650, KU950651, KU950667, KU950670, KU950692, KX655658, KX655662, KX655672, KX765917, KX765920, KX765932, KX765933, KX765941, KX765954, KX765958, KX765971, KX894807, KY460517, KY654507, KY654508, KY654511, KY654512, KY654513, KY654514, KY654518, KY883567, KY967362, KY967364, LC699633, LC699634, LR699737, MF001038, MF001041, MF001047, MF001051, MF001053, MF001054, MF614946, MG642028, MG642030, MG642031, MG642033, MG642048, MG642050, MG642063, MG642070, MG642074, MG773271, MG793382, MH182018, MH182052, MH182061, MH290724, MH447960, MK109775, MK167035, MK749912, MN306017, MN310477, MN531557, MN630093, MN630106, MW020596, MW020597, MW020599, MW582528, MZ515571, MZ515572, MZ515585, MZ515588, MZ515652, MZ515679, MZ515688, MZ515705, MZ515735, MZ515772, MZ515801, MZ515879, MZ515885, MZ515967, MZ516055, MZ516085, MZ516103, MZ516112, MZ516128, OK500256, OK500258, OK500260, OK649589, OK649594, OK649595, OK649596, OK649597, OK649598, OK649600, OK649613, OK649614, OK649625, OK649634, OK649638, OK649642, OK649656, OK649657, OK649658, OK649659, OK649660, OK649661, OK649662, OK649663, OK649668, OK649669, OK649672, OK649674, OK649675, OK649676, OK649677, OK649678, OK649679, OK649680, OK649684, OM857180, OM857196, OM857267, OM857334, ON152648, ON729319, OP744443, OP744445, OP890316, OP890319, OP890322, OP890324, OP890325, OP890327, OP890329, OP890332, OP890337, OP890340, OP965711, OQ024111, OQ024119, OQ024125, OQ024133, OQ024135, OQ024138, OQ024140, OQ024142, OQ024157

*List of accession numbers for reference sequences for RSV-B*

AF013254, AF193331, AY333361, AY353550, AY751111, DQ227364, HQ731688, HQ731708, HQ731711, HQ731714, HQ731718, HQ731722, JF704213, JF704214, JF704224, JN032115, JN032117, JQ582843, JX198143, JX198147, JX198160, JX198165, JX198166, JX489429, JX576730, JX576742, JX576744, JX576746, JX576751, JX576760, JX576761, JX576762, JX976333, JX976378, JX976389, JX976394, KC297428, KC297446, KC297462, KC297466, KC297492, KC342326, KC476971, KF246627, KF246637, KF300963, KF826829, KF826839, KF826843, KF826845, KF826853, KF826860, KJ627262, KJ627285, KJ627302, KJ723476, KJ723481, KJ939919, KJ939926, KJ939928, KJ939929, KJ939932, KM402680, KM402687, KM402697, KM402730, KP258712, KP258713, KP258721, KP258724, KP258731, KP258736, KP258742, KP258745, KP317922, KP317923, KP317928, KP862062, KP862078, KP862288, KP862497, KP862515, KT781406, KU316127, KU316134, KU316173, KU316175, KU316179, KU316181, KU316182, KU950458, KU950467, KU950477, KU950484, KU950588, KU950605, KU950619, KX655648, KX655649, KX655654, KX655669, KX655690, KX765906, KX765912, KX765943, KX765949, KX765957, KX765962, KX775755, KX775765, KX775767, KX775768, KY249657, KY249658, KY249662, KY249670, KY249677, KY249683, KY634397, KY634410, KY684758, KY883571, LC699635, M73545, MF185751, MF185752, MF185754, MF496543, MF496621, MF496628, MG431251, MG431252, MG431253, MG642036, MG642043, MG642062, MG773268, MG813994, MG839547, MN163124, MT040084, MT040085, MT040088, MZ515904, MZ516079, MZ516094, OC699636, OK500262, OK500264, OK649740, OK649745, OK649748, OK649752, OL321917, ON729320, OP699166, OP699167, OP699169, OP699173, OP699177, OP699192, OP699197, OP699202, OP890341, OP890342, OP890343, OP890344, OP890345, OP890346, OP890347, OP890348, OP890349, OP890350, OP965698, OP965700, OP965701, OP965702, OP965703, OP965704, OP965705, OP965706, OP965707, OP965708, OP965710, OQ024159, OQ024160, OQ024161, OQ024162

*List of respiratory syncytial virus sequences generated in this study*

| ***Strain*** | *GenBank Accession* | *GISAID EPI_ISL_* |
|---|---|---|
| hRSV/B/Austria/MUW1119688/2019 | OQ621734 | 16533855 |
| hRSV/A/Austria/MUW1120425/2019 | OQ261746 | 16533868 |
| hRSV/B/Austria/MUW1121202/2019 | OQ261739 | 16533856 |
| hRSV/B/Austria/MUW1123882/2019 | OQ261735 | 16533857 |
| hRSV/B/Austria/MUW1130063/2019 | OQ261736 | 16533858 |
| hRSV/B/Austria/MUW1214013/2020 | OQ261737 | 16533859 |
| hRSV/A/Austria/MUW1228283/2020 | OQ261747 | 16533869 |
| hRSV/A/Austria/MUW1446022/2021 | OQ261748 | 16533870 |
| hRSV/B/Austria/MUW1448780/2021 | OQ261738 | 16533860 |
| hRSV/A/Austria/MUW1451208/2021 | OQ261749 | 16533871 |
| hRSV/A/Austria/MUW1456788/2021 | OQ261750 | 16533872 |
| hRSV/A/Austria/MUW1515402/2022 | OQ261751 | 16533873 |
| hRSV/B/Austria/MUW1554490/2022 | OQ261740 | 16533861 |
| hRSV/B/Austria/MUW1560646/2022 | OQ261741 | 16533862 |
| hRSV/B/Austria/MUW1561030/2022 | OQ261742 | 16533863 |
| hRSV/A/Austria/MUW1561554/2022 | OQ261752 | 16533874 |
| hRSV/B/Austria/MUW1561555/2022 | OQ261743 | 16533864 |
| hRSV/A/Austria/MUW1562468/2022 | OQ261753 | 16533875 |
| hRSV/B/Austria/MUW1563570/2022 | OQ261744 | 16533865 |
| hRSV/B/Austria/MUW1565165/2022 | OQ261745 | 16533867 |

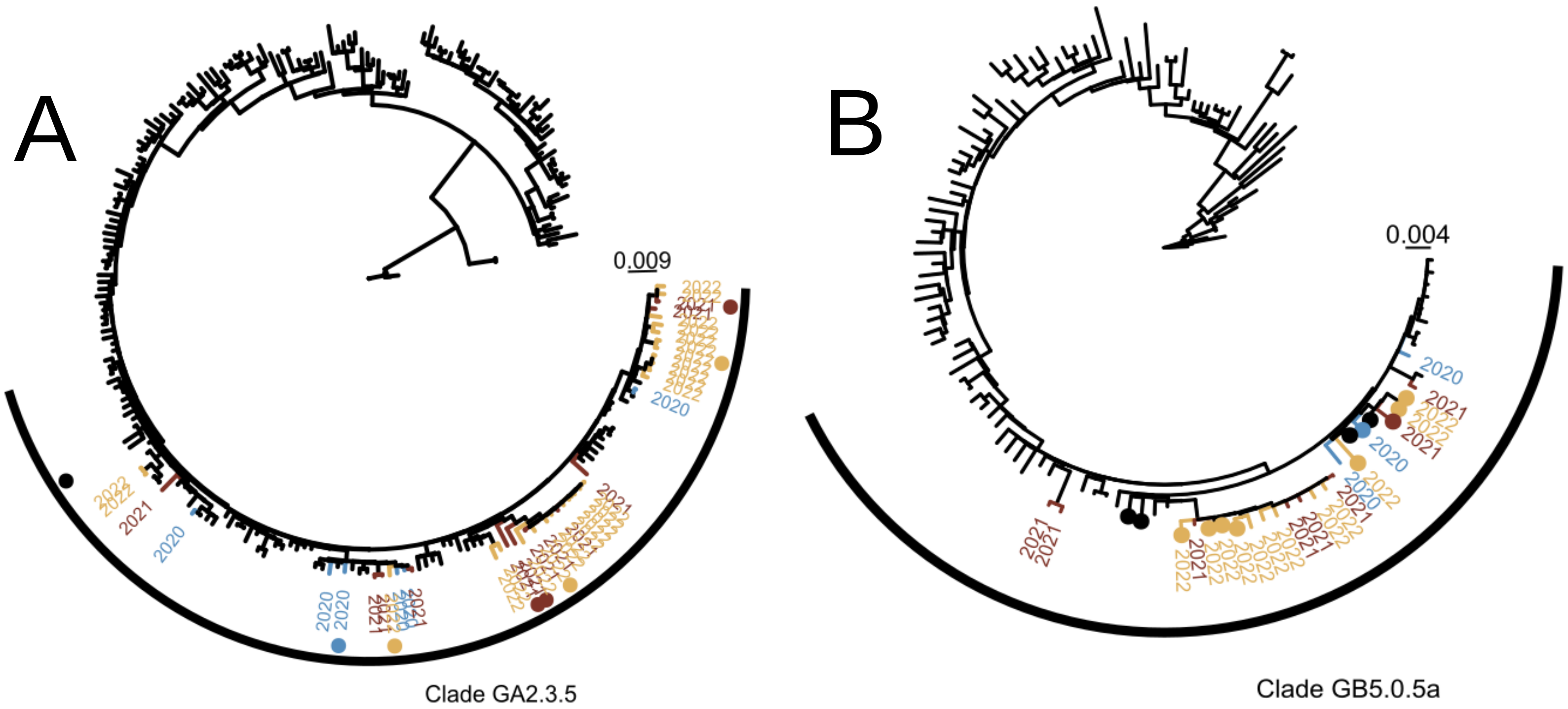


**Figure S1**. **Phylogenetic trees of RSV-A (A) and RSV-B (B) whole genome sequences**. Colored tip labels indicate sequences obtained from 2020 (blue), 2021 (red) and 2022 (yellow); and colored dots indicate sequences obtained from a nationwide sentinel surveillance system in Austria. The trees were inferred over 500 bootstraps using the GTR+G substitution model. Major contemporary clades of RSV-A (GA2.3.5) and RSV-B (GB5.0.5a) are shown by a curved black bar, and the tree scale is shown.

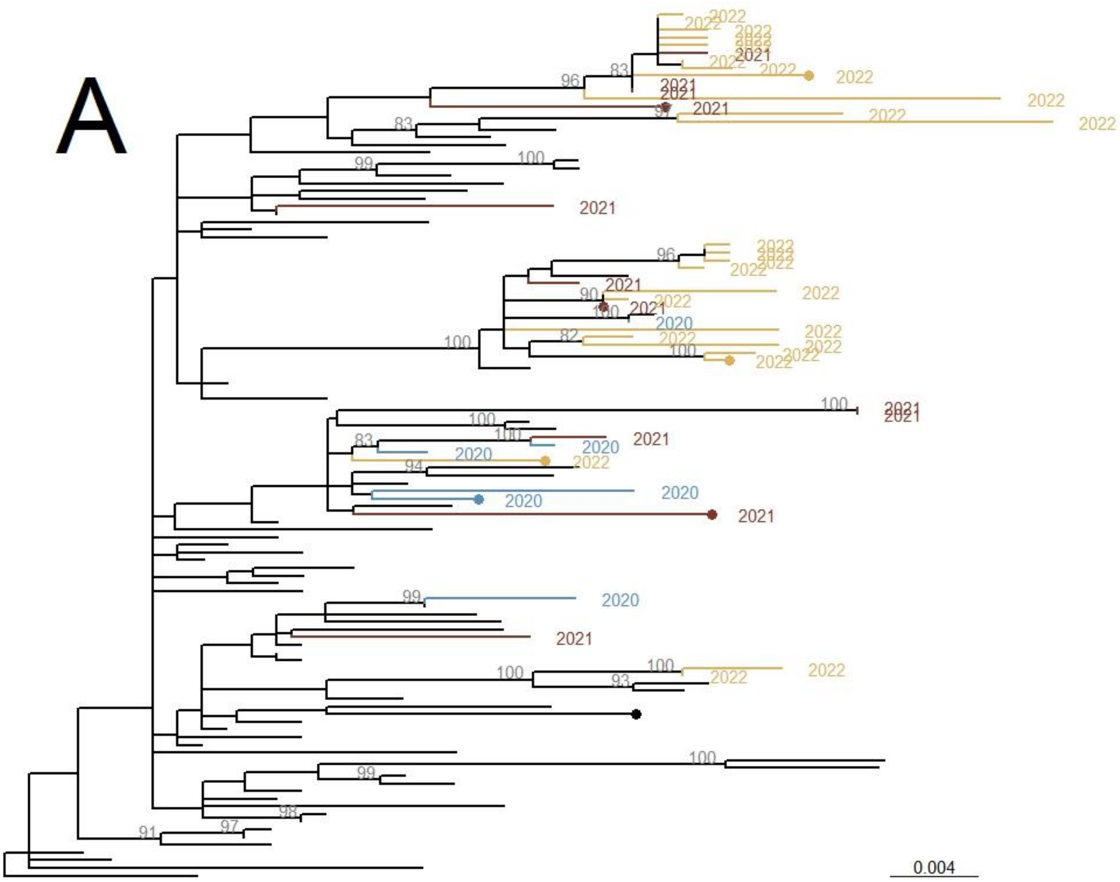


**Supplementary Figure S2.**
Phylogenetic tree of RSV-A based on complete G protein coding sequences, showing only clade GA2.3.5. Colored tip labels indicate year of collection post-COVID-19 pandemic, with colored tip dots indicating sequences from Austria. Bootstrap support is shown in grey beside nodes where ≥80% of trees supported the node. The phylogeny was inferred using the GTR+G substitution model over 500 bootstrap replicates.

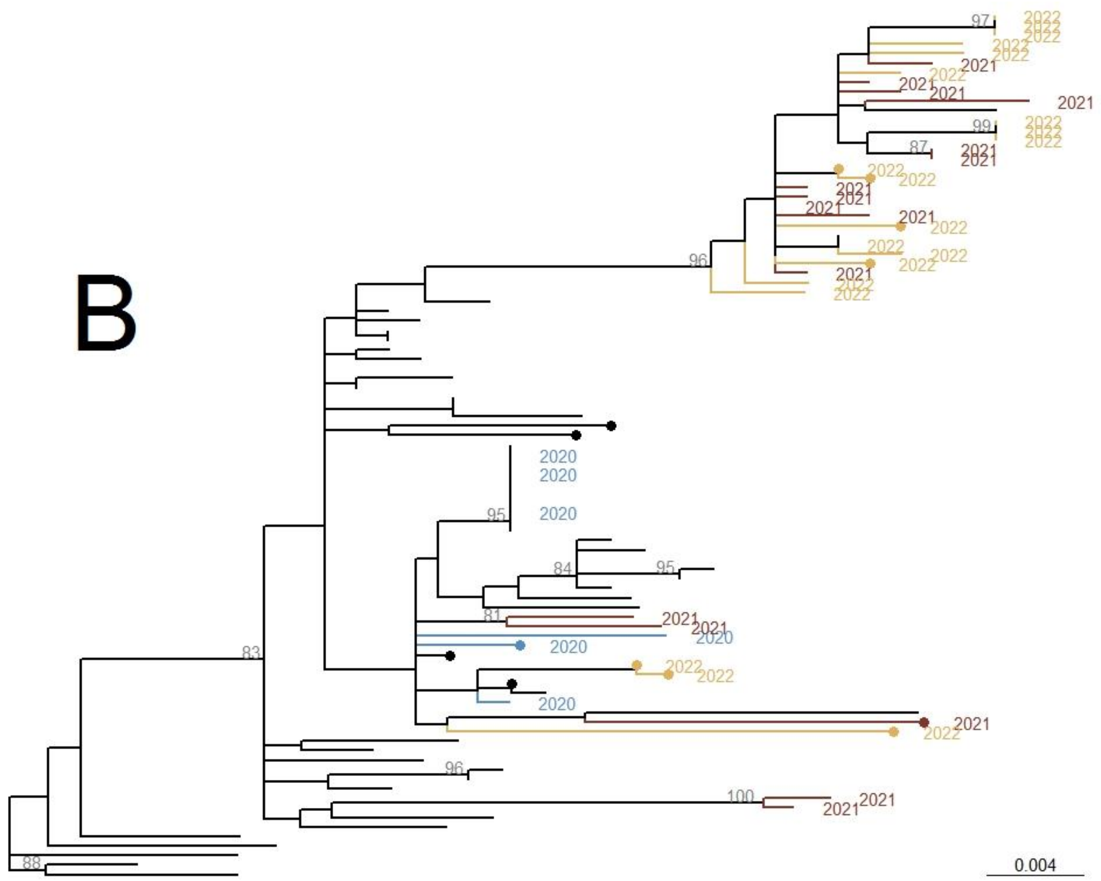


**Supplementary Figure S3.**
Phylogenetic tree of RSV-B based on complete G protein coding sequences, showing only clade GB5.0.5a. Colored tip labels indicate year of collection post-COVID-19 pandemic, with colored tip dots indicating sequences from Austria. Bootstrap support is shown in grey beside nodes where ≥80% of trees supported the node. The phylogeny was inferred using the GTR+G substitution model over 500 bootstrap replicates.
